# The 3’UTR of the *orb2* gene encoding the *Drosophila* CPEB translation factor plays a critical role in spermatogenesis

**DOI:** 10.1101/2020.08.24.264762

**Authors:** Rudolf A. Gilmutdinov, Eugene N. Kozlov, Ludmila V. Olenina, Alexei A. Kotov, Justinn Barr, Mariya V. Zhukova, Paul Schedl, Yulii V. Shidlovskii

## Abstract

CPEB proteins are conserved translation regulators involved in multiple biological processes. One of these proteins in *Drosophila*, Orb2, is a principal player in spermatogenesis. It is required for meiosis and spermatid differentiation. During the later process *orb2* mRNAs and proteins are localized within the developing spermatid. To evaluate the role of *orb2* mRNA 3’UTR in spermatogenesis, we used the CRISPR/Cas9 system to generate a deletion of the *orb2* 3’UTR, *orb2*^*R*^. This deletion disrupts the process of spermatid differentiation, but has no apparent effect on meiosis. While this deletion appears to destabilize the *orb2* mRNA and reduce the levels of Orb2 protein, this is not the primary cause of the differentiation defects. Instead, differentiation appears to be disrupted because *orb2* mRNAs and proteins are not properly localized within the differentiating spermatids. Other transcripts and proteins involved in spermatogenesis are also mislocalized in *orb2*^*R*^ spermatids.

**Author summary:** The conserved family of cytoplasmic polyadenylation element binding (CPEB) proteins can activate or repress translation of target mRNAs, depending on the specific biological context, through interaction with special cytoplasmic polyadenylation element (CPE) sequences. These proteins function mainly in highly polarized cells. Orb2, one of the two *Drosophila melanogaster* CPEB proteins, is predominantly expressed in the testes and is crucial for spermatogenesis. The 3’UTR of *orb2* transcript contains multiple CPE-like motifs, which is indicative of *orb2* self-regulation. We have generated a deletion that removes the greater portion of 3’UTR. While this deletion causes a reduction in the levels of *orb2* mRNA and the protein, this does not appear to be responsible for the defects in spermatogenesis observed in the deletion mutant. Instead, it is the mislocalization of the mRNA and protein in the developing spermatids.

## Introduction

Cytoplasmic polyadenylation element binding (CPEB) proteins are highly conserved translation factors. They interact with special cytoplasmic polyadenylation element (CPE) motifs in the 3’UTRs of target transcripts and can activate or repress their translation depending on the biological context [1, 2]. Vertebrates have four functionally distinct CPEB proteins—CPEB1, CPEB2, CPEB3, and CPEB4— while flies have two, Orb and Orb2. CPEB proteins take part in a broad range of biological processes, including translational control of embryonic cell division [3], cellular senescence [4], and the formation of synaptic plasticity underlying learning and long-term memory [5, 6]. They also have important functions in oogenesis and spermatogenesis. In *Xenopus* oocytes, for example, the sequential activation of CPEB1 and CPEB4 helps to control egg maturation [3, 7–10].

The two CPEB proteins in flies, Orb and Orb2, differ significantly in their N-terminal ends and have largely different activities. Orb is required in the female germline for the proper development of the egg chamber and regulates the translation of oocyte transcripts at multiple time points during this process. It is also expressed during the last stages of spermatogenesis in spermatids and in a subset of mushroom body neurons in the fly brain [11–14]. In contrast, Orb2 is more widely expressed and plays important roles in somatic development and spermatogenesis. During embryonic development, Orb2 is expressed at high levels in the central and peripheral nervous system and functions as a fidelity factor in asymmetric cell division in these tissues and in muscle progenitor cells [15]. In the adult, it has been implicated in learning and memory [6, 16, 17].

In addition to these activities, Orb2 has critical functions during spermatogenesis, and *orb2* mRNA and protein expression levels in adults are the highest in the testes. Two Orb2 protein isoforms are expressed in the testes: the 60-kDa Orb2A isoform and the 75-kDa Orb2B isoform, the latter being more abundant. The two proteins share a 542-amino acid C-terminal sequence that includes several polyQ and polyG amino acid stretches, two RRM-type RNA binding domains, and a zinc finger domain. On the other hand, they have unique N-terminal domains of 162 and 9 amino acids, respectively. The transcripts encoding the larger isoform have relatively long 3’UTRs (580–5791 nt) with multiple CPEs and CPE-like elements, while the transcript encoding the smaller isoform has a short 3’UTR (~400 nt) with no CPEs. The large isoform is essential for male fertility, while the smaller isoform is not [18].

Orb2 expression is not observed at the early stage of spermatogenesis. It is not expressed in the germline stem cells, and only a very low level of this protein can be detected in the mitotic cysts. However, Orb2 expression is substantially upregulated after the 16-cell cysts are formed and the interconnected spermatocytes duplicate their DNA and begin to grow. At this stage as well as during the two meiotic divisions, Orb2 is cytoplasmic and is largely delocalized except puncta around the nuclear envelope [18]. Once the spermatocytes have fully matured, they synchronously enter meiosis I, which is followed by meiosis II. At the end of meiosis, a cyst consisting of 64 interconnected spermatids with haploid nuclei is formed. The cells in the cyst remain undifferentiated during the meiotic divisions but start to differentiate as soon as meiosis is completed. One of the first steps is the reorganization and oriented polarization of the germ cells in the cyst so that all nuclei are clustered towards its basal side (relative to the apical basal polarity of the testes). Basal bodies, which function as microtubule nucleation centers, are located on the apical side of the nuclei. They initiate the assembly of the flagellar axonemes that grow towards the apical tip of the testes. The axonemes elongate until they almost reach this tip and then normally cease to grow [19–22]. During the elongation phase, *orb2* transcript and protein are concentrated in a ring near the tip of the advancing axoneme. Similar to other gene products that are localized during the axoneme growth, *orb2* transcript and protein form a “comet tail” extending back through the axonemes towards the nuclei clustered at the basal pole of the spermatids [18, 23]. Once elongation is complete, the process of individualization begins. Individualization is accomplished by the assembly of a special structure called the individualization complex (IC). The IC consists of 64 actin cones that assemble around the nuclei at the basal cap of spermatid. The IC then travels down the bundled flagellar axonemes, ensheathing each in a plasma membrane and pushing the excess cytoplasm into a waste bag. During this phase, the spermatid nuclei undergo a series of morphological transitions that alter the protein composition of their chromatin [19, 24–26].

Analysis of *orb2* mutants indicates that this gene has important functions in several steps of spermatogenesis [18, 27]. The *orb2* null mutants show no obvious defects in spermatocytes during the prolonged G2 leading up to meiosis I, but these cells are arrested early in meiosis I, accumulating high levels of nuclear cyclin A. In addition to being required in meiosis, Orb2 is also important for spermatid differentiation. In particular, it has a role in the initial polarization of cells in the 64-cell cyst, where this Orb2 appears to have two different functions. One is in orienting cyst polarization relative to the apical– basal axis of the testis. The other is in the polarization of germ cells in the cyst so that their nuclei cluster together at one surface of the cyst, while microtubule nucleation centers (basal bodies) are oriented so that microtubule assembly is directed towards the other (apical) surface of the cyst. Once the 64-cell cyst is properly polarized, *orb2* contributes to the elongation of the flagellar axonemes. It is required for localizing transcripts in a comet pattern in the growing flagellar axonemes and activating their translation. The transcripts whose localization and translation depend on Orb2 include *orb2* and *apkc* mRNAs [18, 27]. *orb2* mutants are also defective in that the growth of the flagellar axonemes is not properly terminated and defective ICs are assembled.

Studies on the other fly CPEB protein, Orb, have shown that it has a positive autoregulatory activity: Orb binds to sequences in the 3’UTR of its own transcript, localizes the transcript to the developing oocyte, and controls its on-site translation. This 3’UTR-dependent autoregulatory activity helps drive oocyte specification in the newly formed 16-cell cysts, while at later stages of oogenesis it is important for ensuring that sufficient levels of Orb protein are localized to the developing oocyte [14, 28]. Similar to the *orb* 3’UTR, most of the *orb2* 3’UTRs are quite long and carry multiple CPE-like elements. Moreover, we and other authors have found that Orb2 is associated with *orb2* transcript *in vivo* [18, 29, 30]. Hence, the question has arisen whether the *orb2* 3’UTR has important functions in spermatogenesis. To address this question, we used the CRISPR/Cas9 system to delete the *orb2* 3’UTR and analyzed the mutants for the effect of this deletion on spermatogenesis.

## Results

### Deletion of the *orb2* 3’UTR

The *orb2* gene is predicted to encode five transcript species that differ in their transcription start sites, splicing patterns, and the lengths of their 3’UTRs. Only one of them, RA, encodes the smaller 60-kDa Orb2A isoform. As shown in Fig 1, RA has a very short 3’UTR that lacks not only the canonical UUUUAU CPE, but also other known CPE-like motifs. All other *orb2* transcripts encode the larger 75-kDa Orb2B isoform. One of them, RB, has a short 580-nt 3’UTR that lacks the canonical UUUUAU CPE, but contains two CPE-like motifs, UUUUUGT and UUUUUGUU, that are reported to be enriched in Orb2-associated mRNAs [30]. The RC transcript has a 1563-nt 3’UTR, while the two remaining transcripts, RD and RH, have 3’UTRs of 3826 and 5791 nt, respectively. The distribution of canonical CPEs in these transcripts is different. There is one canonical and six non-canonical CPEs in the 1563-nt RC 3’UTR, whereas the two larger 3’UTRs have 27 and 37 CPE-like sequences respectively that include 9 canonical UUUUAU motifs.

**Fig. 1.**
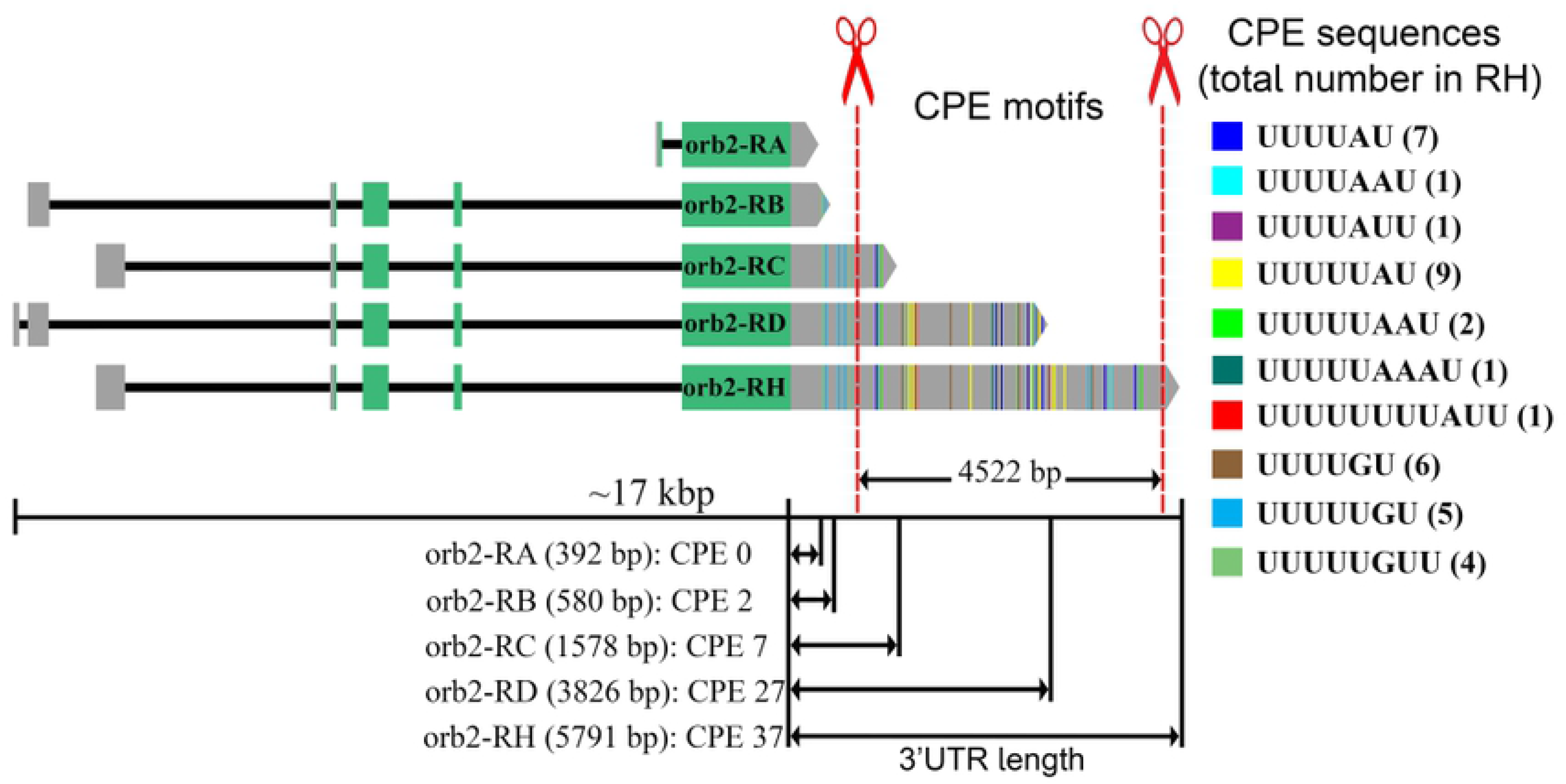
Structure of *orb2* transcripts and location of 3’UTR deletion. The *orb2* gene encodes five distinct mRNA species that differ in the length of their 3’UTRs and contain different sets of canonical and non-canonical CPE sequences (indicated on the right). Cutting sites for CRISPR/Cas9 gene modification are indicated. CPE motifs were taken from the CPEB CLIP dataset [30].

As the RD and RH 3’UTRs contain a much greater number of canonical and non-canonical CPEs, we designed a deletion that selectively removes the bulk of their 3’UTR sequences. As shown in Fig 1, the CRISPR/Cas9 deletion excises a 4522-nt DNA segment that includes all canonical CPEs in RC, RD and RH. Since the proximal endpoint of the deletion is well downstream of the RA and RB polyadenylation signal, these transcripts should not be affected. For RC, the single canonical CPE and one of the non-canonical CPEs are removed together with its predicted polyA addition sequences. The predicted polyadenylation signal of RC and RD transcripts in the deletion is expected to be the same as that of RH. For all three of these transcripts, the 3’UTR sequence upstream of the deletion breakpoint is 1008 nt long and contains five non-canonical but Orb2-enriched CPEs (UUUUUGT or UUUUUGUU).

In the initial fly stock, the deleted DNA was replaced by the *DsRed* gene flanked by loxP sites and an attP sequence. *DsRed* was then excised to give the *orb2* gene carrying attP and loxP sites in place of the 4522-nt deletion. The deletion was verified by sequencing, and the resulting mutation was designated *orb2*^*R*^.

### Most *orb2*^*R*^ males are sterile

While *orb2* null alleles are semi-lethal, with only a few flies surviving to adulthood, the *orb2*^*R*^ mutation does not appear to affect any essential processes during development, since the number of homozygous *orb2*^*R*^ flies reaching the adult stage is close to that in wild-type (WT) flies (Suppl. Fig 1). On the other hand, male fertility in the *orb2*^*R*^ mutants is reduced. Figure 2 shows the results of experiment where 35 *orb2*^*R*^ males were mated with 70 WT females. After 10-day incubation, adult flies were removed from the vials and their offspring were allowed to develop to adulthood. Quantification of the number of offspring indicates that the overall fertility of *orb2*^*R*^ males is reduced approximately tenfold. Two scenarios could potentially explain the reduction of male fertility in *orb2*^*R*^ flies. First, the production of functional sperm could be more or less uniformly impaired in all *orb2*^*R*^ males. An alternative, though seemingly less likely possibility is that the fertility of individual flies could be affected differentially, so that some males are fertile while others are sterile. To distinguish between these alternatives, we measured the fertility of individual *orb2*^*R*^ males. In the first experiment, we mated individual males to two WT virgin females for one week and then scored the number of males that produced offspring. Unexpectedly, we found that second scenario was correct: about 75% of the *orb2*^*R*^ males were completely sterile (Fig. 2B). When *orb2*^*R*^ was placed in *trans* to the null allele *orb2*^*36*^ (a deletion of *orb2*), no offspring were produced. Moreover, the number of offspring from the few fertile *orb2*^*R*^ males is substantially reduced, compared to WT or to males heterozygous for *orb2*^*36*^. In the experiment shown in Fig. 2C, we mated males of the above genotypes to WT females and then scored the number of offspring they produced. WT males typically have more than 60 offspring, while males heterozygous for an *orb2* null allele, *orb2*^*36*^, have slightly fewer, with an average of a bit more than 60 offspring. In contrast, of the fertile *orb2*^*R*^ males most had substantially reduced fertility and produced fewer than 30 offspring.

**Fig. 2.**
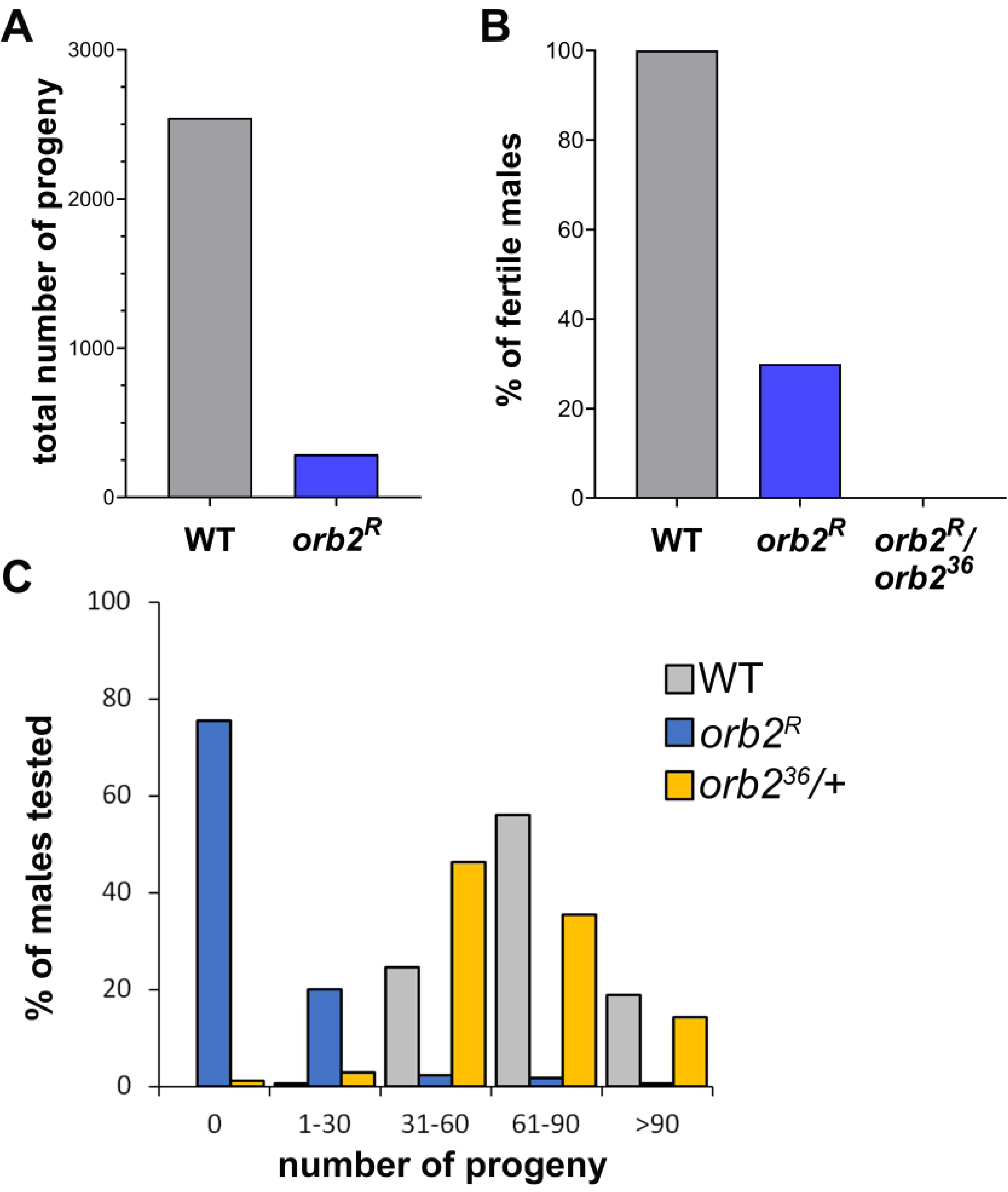
Fertility assay. **(A)** Overall fertility: 35 WT or *orb2*^*R*^ males were mated with 70 WT females, and the total number of offspring reaching adulthood was counted. **(B)** Individual fertility: males of the WT, *orb2*^*R*^ and *orb2*^*R*^/*orb2*^*36*^ genotypes were each mated with two WT virgin females for 7 days, and the number of males that produced any offspring was counted (WT, *n* = 287; *orb2*^*R*^, *n* = 358; *orb2*^*R*^/*orb2*^*36*^, *n* = 145). **(C)** Frequency distribution of offspring from individually mated males. Males of the WT, *orb2*^*R*^ and *orb2*^*36*^/+ genotypes were crossed for 7 days with WT virgin females. The crossed flies were then removed, and the numbers of offspring from each individual male were estimated and ranged into groups. All data are represented as percentage relative to the total number of males tested. Tests were conducted with 175 males of each genotype.

### The accumulation of *orb2* transcript and protein is reduced in *orb2*^*R*^ testes

To better understand the nature of spermatogenesis defects in the *orb2*^*R*^ mutant, we examined the expression of both *orb2* mRNA and Orb2 protein. Quantitative RT–PCR (reverse transcription– polymerase chain reaction) was used to measure the relative levels of *orb2* transcript in the testes of WT and *orb2*^*R*^ males. The *GADPH* transcript, which lacks canonical CPEs, served as a control for RNA input. As shown in Fig. 3A, the level of *orb2* mRNA in *orb2*^*R*^ testes is reduced approximately by half, compared to WT. This appears to be due to the decreased stability of the mRNA, since the level of *orb2* primary transcripts (detected with primers located on the intron–exon junction) in *orb2*^*R*^ testes is close to that in WT.

**Fig. 3.**
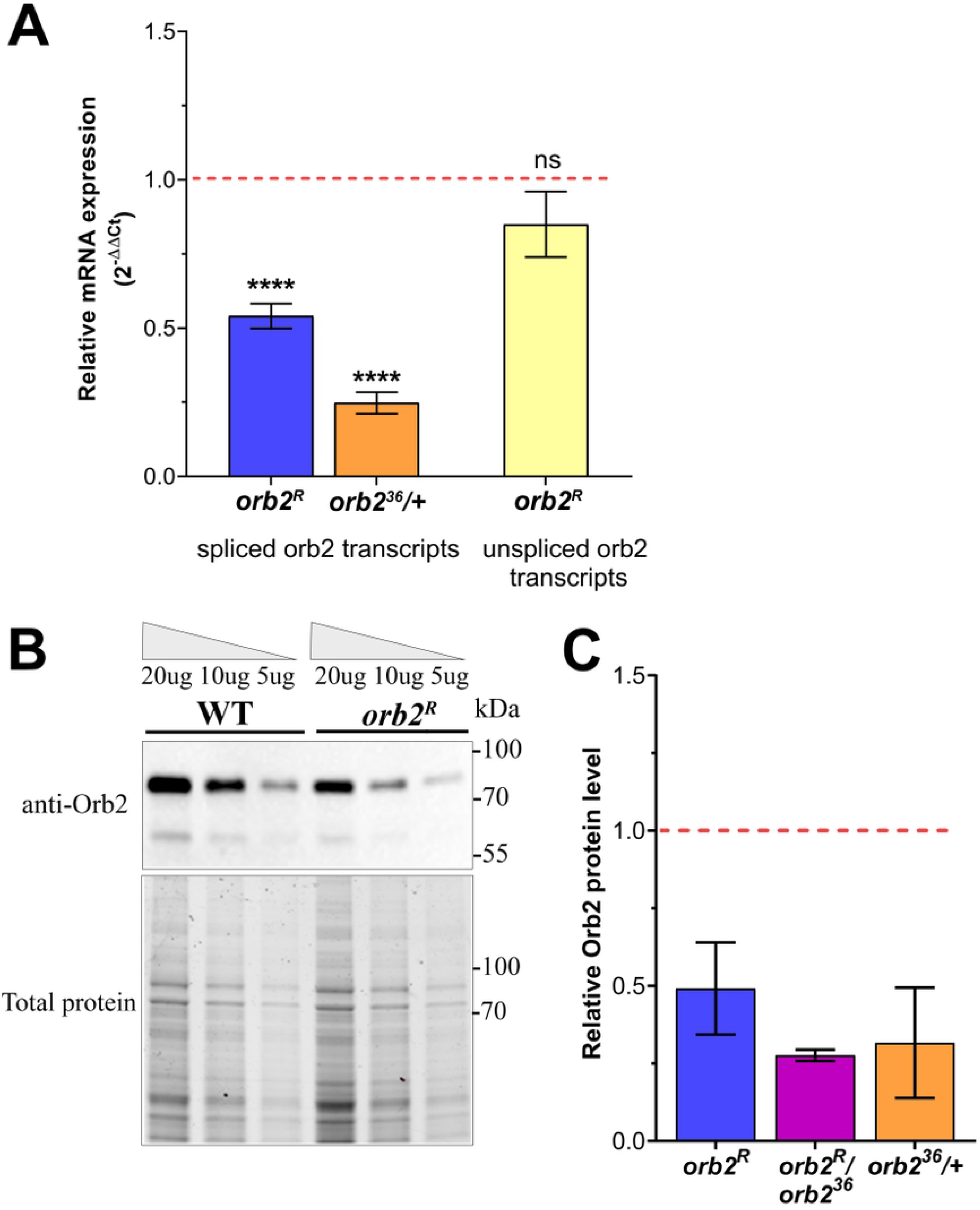
Deletion in 3’UTR of *orb2* reduces its expression in testes. **(A)** The level of total *orb2* mRNA in *orb2*^*R*^ and *orb2*^*36*^/+ mutant testes and the level of primary *orb2* transcripts in *orb2*^*R*^ mutant testes. Data were first normalized to the expression of *GAPDH* in testes and expressed as the mean of fold change [2^−ΔΔCt^] ± SEM in mutant testes relative to control ones (red dashed line) (spliced isoform: control, *n* = 29; *orb2*^*R*^, *n* = 27; *orb2*^*36*^/+, *n* = 18; unspliced isoform: control, *n* = 10; *orb2*^*R*^, *n* = 10). Unpaired two-tailed t-test: **** *P* < 0.0001; ns, *P* > 0.05. **(B)** Western blot analysis of Orb2 protein in twofold dilution series of testes lysates (total protein level is indicated above). A fragment of gel stained for total protein is shown below. **(C)** Densitometry analysis of Orb2 level normalized to total protein level (Bio-Rad stain-free technology) in mutant testes compared with WT testes (red dotted line). Quantitative data of western blotting were obtained from independent biological replicates (*orb2*^*R*^, *n* = 7; *orb2*^*R*^/*orb2*^*36*^, *n* = 4) and expressed as mean ± SD.

Both Orb2 isoforms were detected in Western blots of extracts from *orb2*^*R*^ testes, but the 75-kDa isoform in mutants proved to be reduced, compared to WT. The difference in the levels of this protein is illustrated in the blot of serial dilutions of the extracts from WT and *orb2*^*R*^ males (Fig. 3B). Quantitative analysis (Fig. 3C) showed that the 75-kDa isoform of Orb2 in mutant males was reduced approximately twofold, as in the case of *orb2* mRNA.

The above experiments indicate that the deletion of sequences in the 3’UTRs of the *orb2* transcripts results in a twofold reduction in both mRNA and protein levels, while transcription is unaffected. This finding suggests that the deletion mutant mRNAs are less stable. However, it is surprising that a two-fold reduction in the level of mRNA and protein is sufficient to significantly perturb spermatogenesis so that most *orb2*^*R*^ males were sterile. In fact, there is no evidence of a strong haploinsufficiency as males heterozygous for the *orb2* deletion (*orb2*^*36*^) produced nearly as many offspring as WT males (Fig. 2C) and showed no obvious abnormalities in spermatogenesis. To confirm that *orb2*^*36*^/+ testes have the expected two-fold reduction in *orb2* gene products, we compared *orb2* mRNA and protein levels in WT and *orb2*^*36*^/+. As shown in Fig. 3A, the amount of *orb2* mRNA in *orb2*^*36*^/+ testes was only about a quarter that in WT. By contrasts, *orb2* mRNA was only reduced about two-fold in *orb2*^*R*^. Similar results were obtained when we compared protein levels in WT and either *orb2*^*36*^/+ or *orb2*^*R*^/*orb2*^*36*^. The level of Orb2 protein in *orb2*^*36*^/+ was about one third that in WT, and about one quarter that in *orb2*^*R*^/*orb2*^*36*^.

These findings indicate that the reduction in *orb2* mRNA and protein in homozygous *orb2*^*R*^ testes is if anything less than that in *orb2*^*36*^ heterozygotes, while the fertility of these flies is significantly different. Hence, it is unlikely that the reduction in mRNA and protein in *orb2*^*R*^ testes is in itself responsible for the significantly reduced fertility of the 3’UTR deletion mutant.

### The *orb2* 3’UTR is required for proper *orb2* transcript and protein localization

To better understand why most *orb2*^*R*^ males are sterile, we examined *orb2* transcript and protein expression during spermatogenesis. The pattern of their accumulation in premeiotic and meiotic cysts is similar to that in WT. The expression of *orb2* transcript and protein in *orb2*^*R*^ testes is upregulated after the formation of the 16-cell spermatocyte cysts, and the protein is distributed more or less evenly throughout the cytoplasm. However, the amounts of transcript and protein are reduced, compared to WT (Suppl. Fig. 2). Unlike in flies homozygous for the null-allele *orb2*^*36*^, spermatogenesis in *orb2*^*R*^ does not arrest prior to meiosis I even though Orb2 protein level are lower than in WT; instead, both meiotic divisions appear to be normal, and 64-cell cysts are formed.

While spermatogenesis appears to be unaffected through the completion of meiosis, a series of abnormalities become evident once the spermatids begin to differentiate. When they are first formed, the 64-cell cysts in *orb2*^*R*^ resembled those in WT. The haploid cells in the cyst have a round shape and are about ~10 μm in diameter. Subsequently, the cyst begins to polarize. All nuclei cluster towards the basal side of the cyst, while the basal body associated with each nucleus localizes to the apical side of the cysts and initiates the assembly of the flagellar axoneme. The flagellar axonemes then begin elongating towards the apical end of the testis and ultimately form elongated cells that are almost 2 mm long [19, 21, 22].

During elongation, *orb2* transcript and protein concentrate in a band near the growing tip of flagellar axoneme, with a comet tail extending back towards the nuclei. This is illustrated in Figs. 4A and 4B, respectively, and the distribution of *orb2* transcript and protein along the flagellar axoneme is quantified in Fig. 4C. A different result is obtained in *orb2*^*R*^ mutant testes, or when *orb2*^*R*^ is in *trans* to the *orb2*^*36*^ deletion. Instead of accumulating near the tip of the flagellar axonemes, *orb2*^*R*^ transcripts in *orb2*^*R*^/*orb2*^*R*^ and *orb2*^*R*^/*orb2*^*36*^ mutant testes are concentrated mainly in the middle regions of the axoneme, while their level near the top is substantially reduced (Figs. 4A, 4C). Likewise, the amount of Orb2 protein in WT also increases near the tip (Fig. 4C), but this is not the case in *orb2*^*R*^ or in *orb2*^*R*^/*orb2*^*36*^ (see Figs. 4B, 4C). These findings indicated that the deleted 3’UTR sequences are important for the proper localization of *orb2* transcript and protein during flagellar axoneme elongation.

**Fig. 4.**
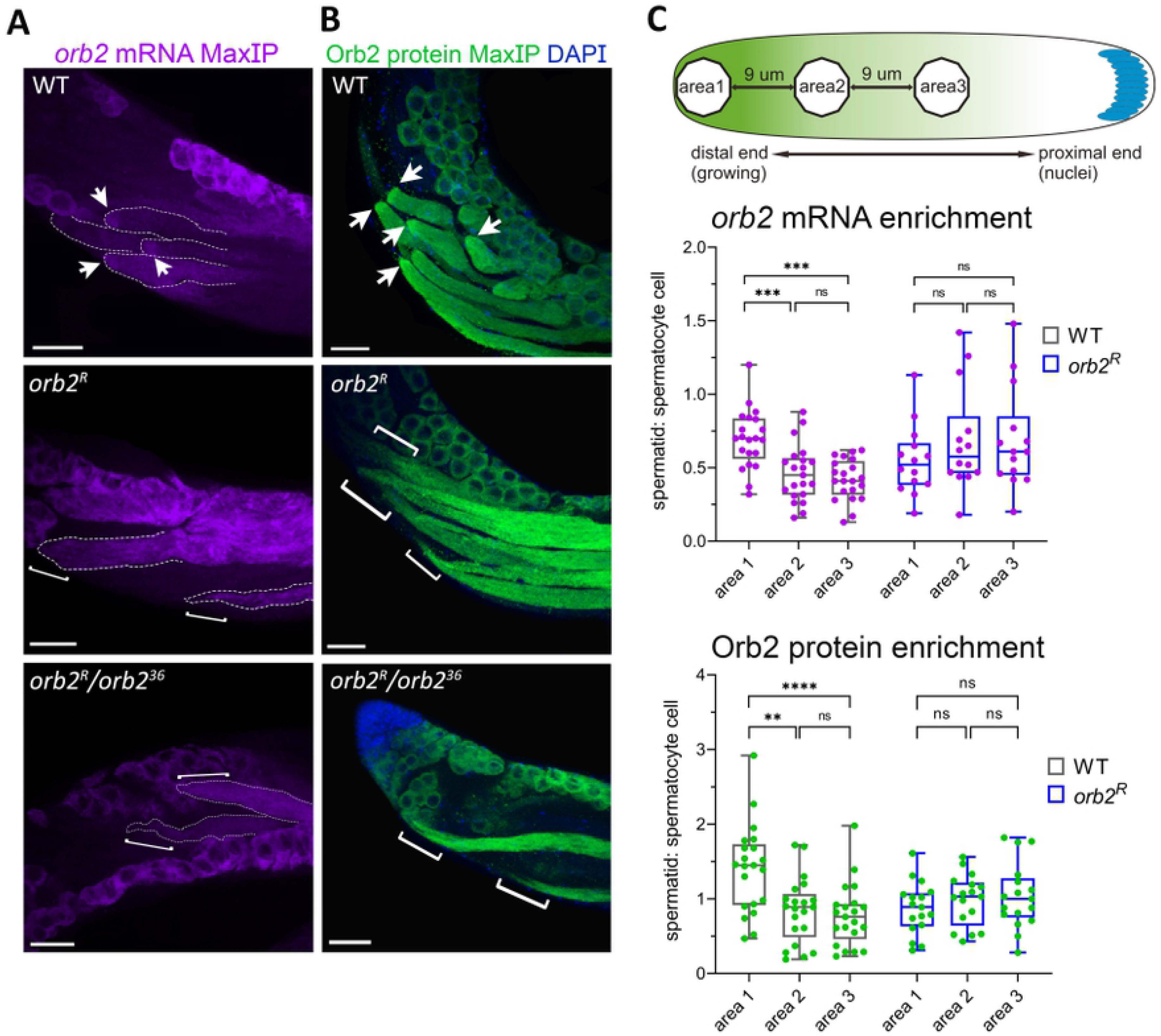
The *orb2* 3’UTR is required for localization of mRNA and protein in spermatid cysts. **(A, B)** Maximum intensity projections of (A) *orb2* mRNA and (B) Orb2 protein within spermatid cysts in WT, *orb2^R^,* and *orb2*^*R*^/*orb2*^*36*^ testes. Arrows indicate the mRNA or protein accumulated at the ends of WT spermatid cysts; brackets indicate the spermatid cyst tail ends without corresponding mRNA or protein accumulation in mutant testes. Scale bar, 40 μm. **(C)** Quantification of *orb2* mRNA and protein distribution along the spermatid cyst. Areas analyzed within a spermatid cyst are shown at the top. Below are box plots of *orb2* mRNA and protein levels in the different areas of spermatid cysts relative to those in spermatocytes for WT (*n* = 21) and *orb2*^*R*^ (*n* = 14). **** *P* < 0.0001, *** *P* < 0.0005, ** *P* < 0.005; ns, not significant.

### The *orb2*^*R*^ mutation affects the axonemal localization of other transcripts and proteins

A number of other transcripts and proteins have been found to have a comet-like distribution in elongating flagellar axonemes [18, 23]. One of these is *orb*, which encodes the other fly CPEB protein. In WT *orb* mRNA preferentially accumulates in a band near the tip of the elongating flagellar axonemes (see Fig. 5A); however, it does not appear to be translated until the late elongation phase, when Orb2 protein begins to disappear. Abnormalities in the localization of *orb* transcript and the expression of Orb protein were evident in *orb2*^*R*^ and *orb2*^*R*^/*orb2*^*36*^ testes. Instead of being localized to the tip of the elongating axonemes, *orb* transcript in *orb2*^*R*^ was distributed over much of the axoneme (Fig. 5A). In addition to being delocalized in *orb2*^*R*^/*orb2*^*36*^ testes, the levels of *orb* transcript in the axonemes are also reduced. In line with the disruption in transcript localization, Orb proteins did not show preferential accumulation at the tip of the flagellar axonemes, and also were present prematurely. As indicated by the brackets in Fig 5B, the tips of the flagellar axonemes in *orb2*^*R*^ and *orb2*^*R*^/*orb2*^*36*^ testes contained little Orb protein. Instead, Orb either accumulated in an intermediate position or was distributed over much of the flagellar axonemes.

**Fig. 5.**
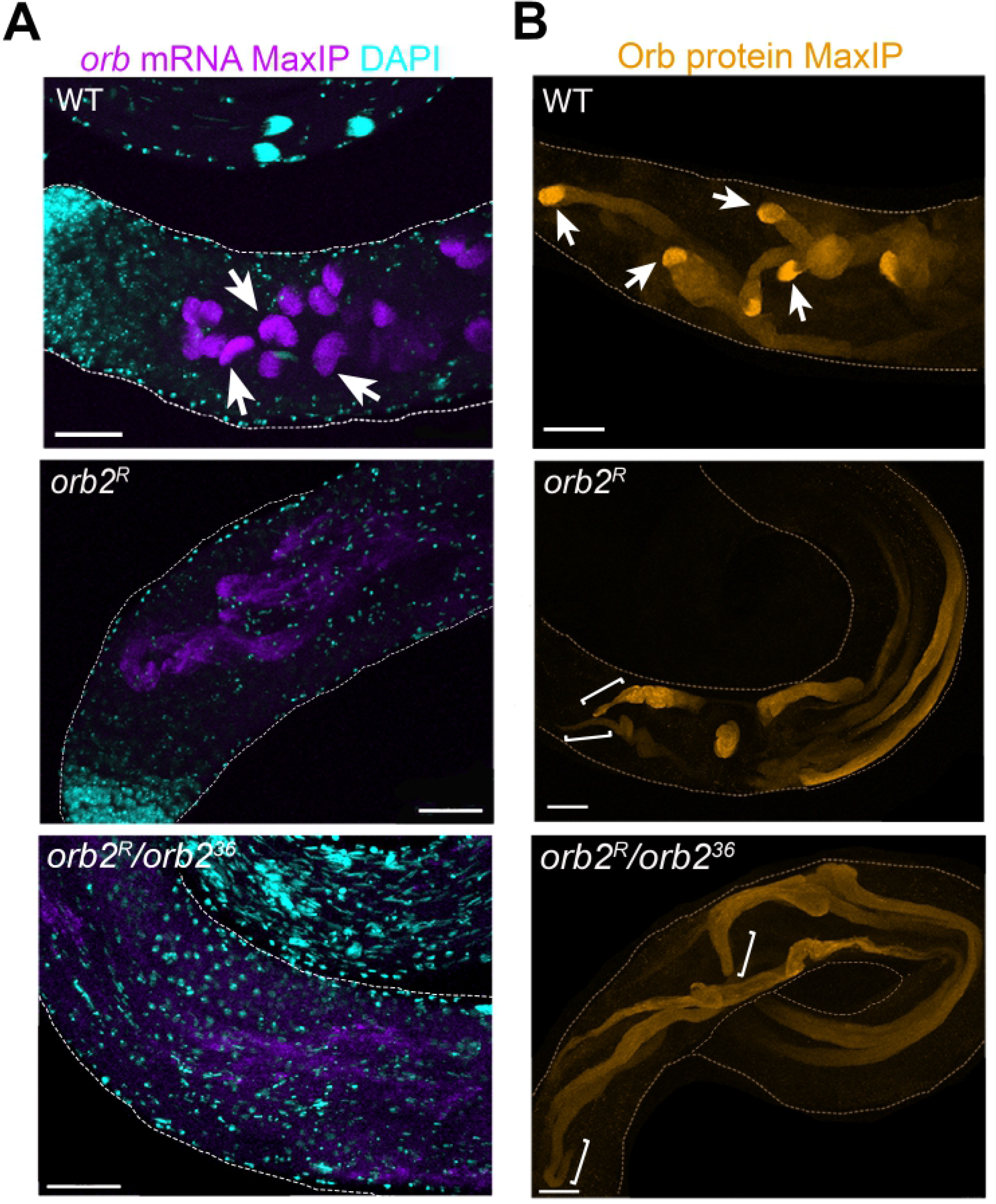
*orb* mRNA and protein localization in testes depend on *orb2* 3’UTR. **(A)** Maximum intensity projections of *orb* mRNA in WT (*n* = 15), *orb2*^*R*^ (*n* = 24), and *orb2*^*R*^/*orb2*^*36*^ (*n* = 20) testes. A high fluorescence signal is observed at the ends of WT spermatid tails (arrows), whereas this pattern in mutant testes is lost, and *orb* mRNA is distributed uniformly. **(B)** Maximum intensity projections of Orb protein in WT (*n* = 30), *orb2*^*R*^ (*n* = 28), and *orb2*^*R*^/*orb2*^*36*^ (*n* = 13) testes. Arrows indicate Orb protein localization at the ends of WT spermatid cysts; brackets indicate the spermatid cyst tail ends without Orb protein accumulation in mutant testes. Scale bar, 50 μm.

The fly homolog of the mammalian DAZ fertility factor is the RNA binding protein Boule (Bol) [31]. The Bol protein can be co-immunoprecipitated with Orb2 in an RNase-resistant complex, and during the spermatid elongation phase it co-localizes with Orb2 in a region near the tip of the growing flagellar axonemes [18]. There is also a comet-like gradient that extends back from the tip towards the nuclei on the basal side of spermatids (Fig. 6). This pattern of localization is not observed in *orb2*^*R*^ or *orb2*^*R*^/*orb2*^*36*^ testes. Unlike in WT, Bol is not preferentially localized close to the end of elongating axoneme (see brackets in Fig. 6). Instead, it is distributed more or less uniformly over much of the axoneme.

**Fig. 6.**
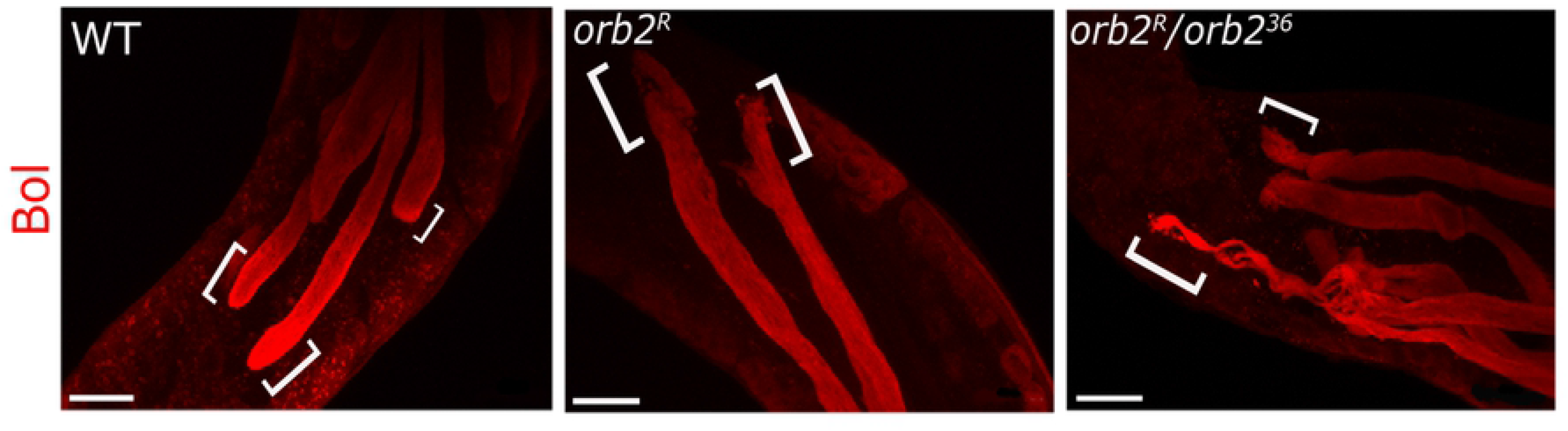
The *orb2* 3’UTR deletion affects the Boule protein localization. Maximum intensity projections of Boule protein in WT and mutant testes. Boule is enriched at the ends of elongated spermatid cysts in WT (*n* = 9) but is uniformly distributed along spermatid cysts in *orb2*^*R*^ (*n* = 22) and *orb2*^*R*^/*orb2*^*36*^ (*n* = 32). Brackets indicate the spermatid cyst tail ends. Scale bar, 40 μm.

### Organization of nuclei in early and late *orb2*^*R*^ spermatids

The polarization of the 64-cell cysts is one of the first steps in spermatid differentiation. The nuclei cluster towards the basal side of the cyst, while the basal bodies associated with each nucleus anchor the flagellar axonemes on the apical side of the cyst [19, 21, 22]. While *orb2*^*R*^ and *orb2*^*R*^/*orb2*^*36*^ cysts assemble flagellar axonemes, there are defects in the initial clustering of nuclei on the basal side of the cyst, and the nuclei are found randomly distributed in the elongating flagellar axonemes (see Fig. 7). As the spermatid tails grow, the nuclei in WT undergo a series of morphological changes. Initially they have a spherical shape but then undergo transition through several intermediate stages, including the leaf, early canoe, late canoe, and finally needle stage [19, 21, 22]. Along with these morphological changes, the nuclei coalesce into a tight bundle to form an inverted cap-like structure (Fig. 8A) [22]. In *orb2*^*R*^ testes, most of the nuclei in the cysts appear to progress to the needle stage, but their subsequent coalescence into the cap-like structure is defective, with only a few exceptions (~10% of the testes) (Figs 8A, 8C). In about 45% of the testes, only a subset of the spermatid cysts has nuclei that coalesced into a cap-like structure, while in other cyst the nuclei are scattered or display only partial coalescence. No coalesced nuclei were found in the remaining testes (~45%) (Fig. 8C). When *orb2*^*R*^ is *trans* to *orb2*^*36*^, only cysts with scattered nuclei are observed.

**Fig. 7.**
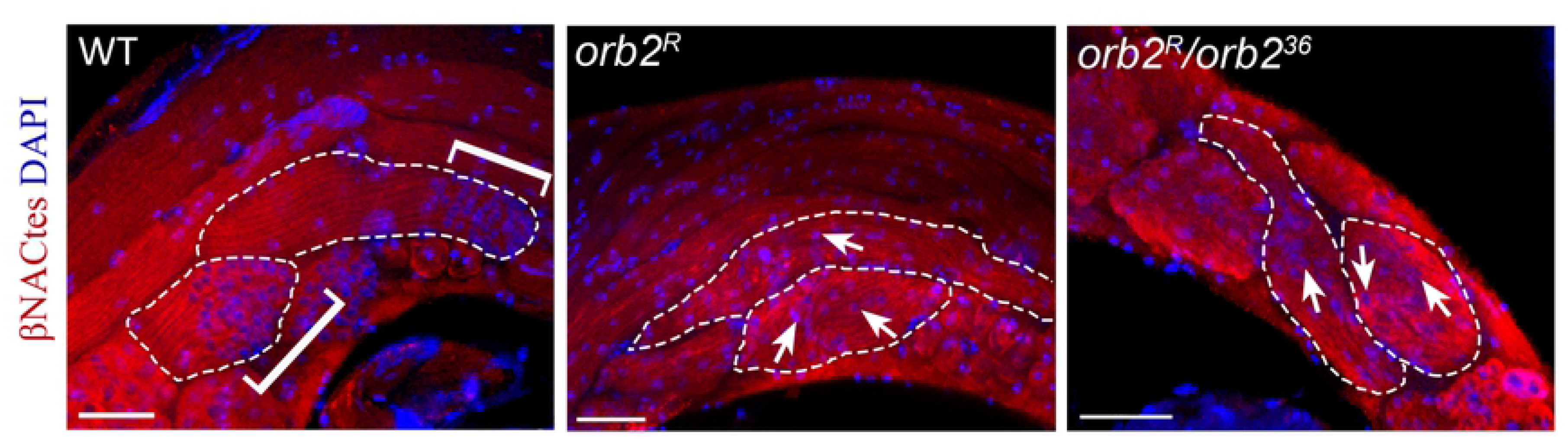
*orb2* 3’UTR is required for nuclear polarization in early elongated spermatid. Whole mount testis staining with βNACtes antibodies (red), which mark germline cells in testes [43]. Chromatin was stained by DAPI (blue). Brackets indicate the areas of nuclear polarization at the proximal ends of early elongated spermatid cysts in WT (*n* = 29). In contrast, the distribution of nuclei (arrows) along elongated spermatid is uniform in *orb2*^*R*^ (*n* = 34) and *orb2*^*R*^/*orb2*^*36*^ (*n* = 36). Scale bar, 30 μm.

**Fig. 8.**
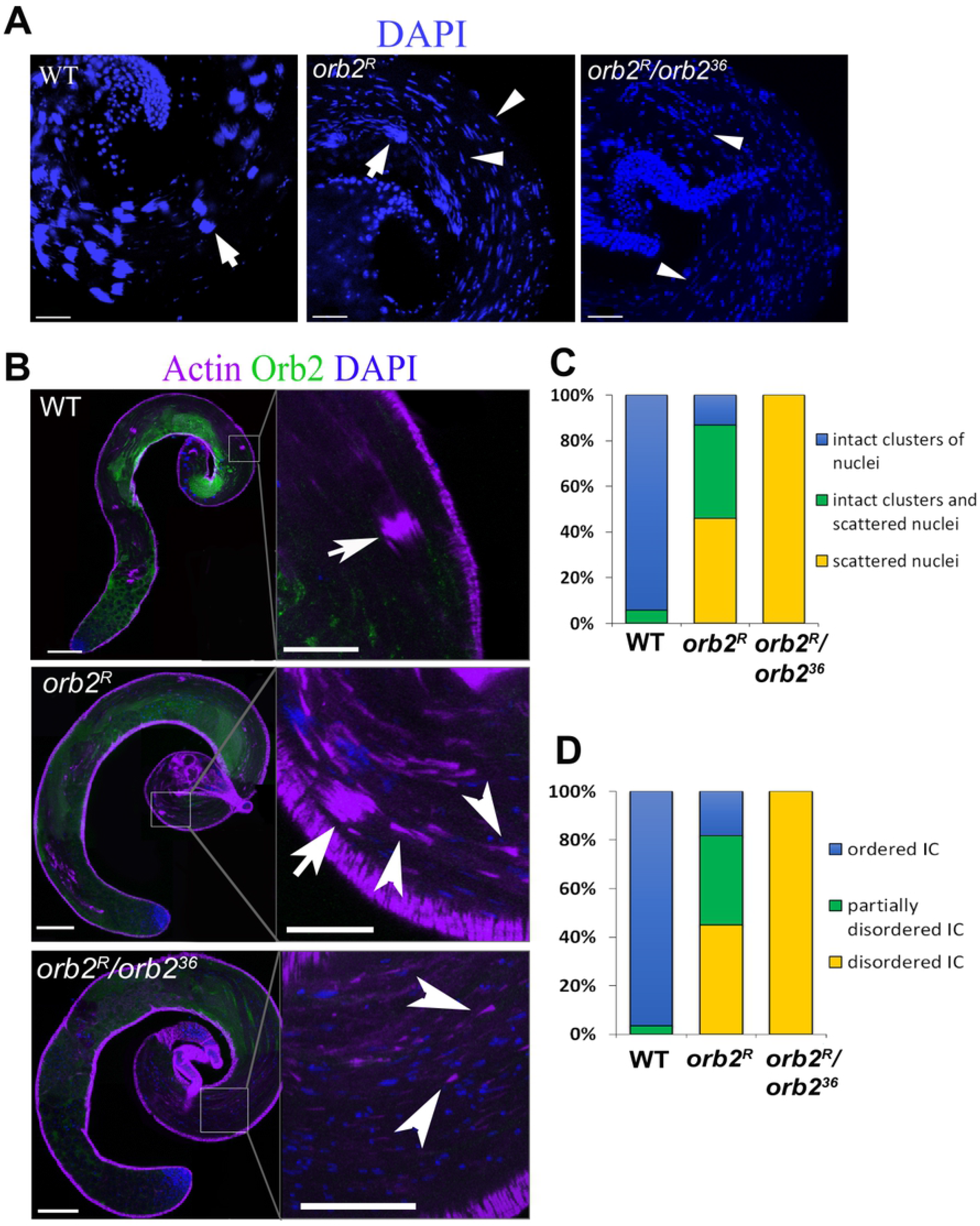
The compaction of nuclei and formation of IC is disrupted in late *orb2*^*R*^ spermatids. **(A)** Confocal slices of immunostained whole mount testis preparations are shown. Chromatin was stained by DAPI (blue). The arrow indicates a condensed spermatid nuclear bundle in WT and partially assembled spermatid nuclear bundles in *orb2*^*R*^. Arrowheads indicate scattered nuclei incapable of compaction in mutant spermatids. **(B)** Individualization complexes are not properly assembled in *orb2*^*R*^and *orb2*^*R*^/*orb2*^*36*^. Confocal slices of whole mount testis preparations are shown. Chromatin was stained by DAPI (blue), actin cones were stained using phalloidin (violet). The arrow indicates complete ICs in WT and incomplete ICs in *orb2*^*R*^. Arrowheads indicate scattered actin cones in mutant testes. Scale bar: left panels, 100 μm; right panels, 30 μm. **(C)** The frequency of spermatids defective in nuclei clustering in WT (*n* = 86), *orb2*^*R*^ (*n* = 61), and *orb2*^*R*^/*orb2*^*36*^ (*n* = 57). **(D)** Quantification of the numbers of testes with IC defect in WT (*n* = 86), *orb2*^*R*^ (*n* = 60) and *orb2*^*R*^/*orb2*^*36*^ (*n* = 56).

### Individualization complex was not properly assembled in mutants

When flagellar axoneme elongation is complete, the spermatids enter the individualization stage. The actin-rich individualization complex (IC) is assembled around each nucleus in a cap-like structure. The IC then begins to move down the flagellar axonemes, investing each spermatid with its own plasma membrane and extruding the excess cytoplasm into a “waste bag” [24, 32]. ICs are successfully assembled in only about 15% of the *orb2*^*R*^ testes (Figs 8B, 8D) In the remaining *orb2*^*R*^ testes, either only a subset of the elongated spermatids assemble an IC or there is no IC assembly at all. IC assembly in *orb2*^*R*^/*orb2*^*36*^ *trans*-heterozygotes is completely disrupted.

Consistent with the defects in the assembly of ICs, only about 30% of seminal vesicles in *orb2*^*R*^ are filled with mature sperm, while others are either empty or filled only partially (Fig. 9). Seminal vesicles in *orb2*^*R*^/*orb2*^*36*^ contain no functional sperm. These findings are consistent with data on the fertility of *orb2*^*R*^ and *orb2*^*R*^/*orb2*^*36*^ males.

**Fig. 9.**
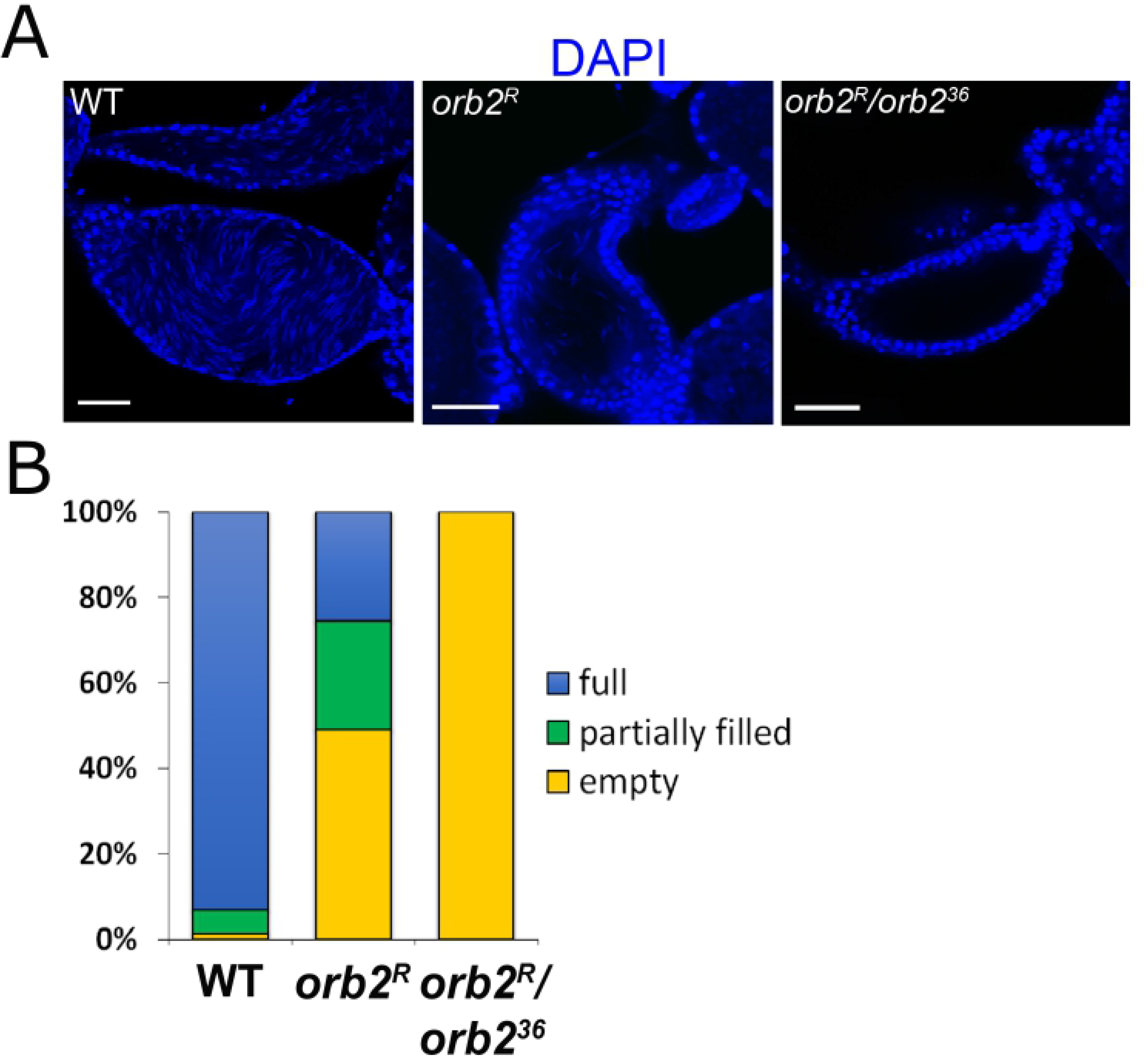
Defects of seminal vesicle filling in *orb2*^*R*^ and *orb2*^*R*^/*orb2*^*36*^. **(A)** Staining for nuclei by DAPI shows that seminal vesicles in *orb2*^*R*^ are filled only partially, while those in *orb2*^*R*^/*orb2*^*36*^ are empty. Scale bar, 40 μm. **(B)** Proportions (%) of seminal vesicles with filling defects in WT (*n* = 74), *orb2*^*R*^ (*n* = 47), and *orb2*^*R*^/*orb2*^*36*^ (*n* = 37).

## Discussion

The localization of gene products to the cellular domains where their functions are required is critical to the establishment of cell polarity. Depending on the context, a variety of mechanisms can be employed to ensure proper targeting [33, 34]. One of them involves the on-site translation of localized transcripts. After their synthesis and export, the translationally silenced transcripts are localized either by an active microtubule-dependent mechanism or by passive diffusion. Once the transcripts are on site, RNA-binding proteins interact with them to regulate their translation. The CPEB protein family is a group of translation factors that help anchor and control the on-site translation of localized transcripts [35–40]. CPEBs recognize CPE elements in the 3’UTR of localized transcripts and can function to repress or activate their translation, depending on the context. The canonical CPE sequence is UUUUAU; however, several variants of this motif are enriched in transcripts that are found associated with different members of the CPEB family. *Drosophila* has two CPEB proteins, Orb and Orb2. The former has essential functions during oogenesis and is required for the translation of multiple oocyte-localized transcripts [12, 13, 41]. Moreover, it also has a autoregulatory activity, with Orb binding to the *orb* transcript 3’UTR and activating its own expression [14]. This autoregulatory activity plays a key role in oocyte specification, and *orb* mutants that lack portions of the *orb* transcript 3’UTR fail to specify an oocyte [28].

Although *orb2* has no essential function in oogenesis, it is required at several stages of spermatogenesis [18, 27]. Here, we have investigated the role of *orb2* 3’UTR sequences in the transcripts encoding the larger 75-kDs isoform in *orb2* activity during spermatogenesis. Four transcripts (RB, RC, RD, and RH) are predicted to encode the 75-kDa isoform. They carry 3’UTRs of different lengths with different numbers of CPE and CPE-like elements, from 2 (RB) to 37 (RH). We generated a deletion that removed 32 out of 37 CPE-like elements, and the resultant allele was named *orb2*^*R*^. This deletion was downstream of the RB polyadenylation site but included the RC and RD polyadenylation sites. As a result, RC, RD, and RH were predicted to all have the same polyadenylation site and contain a total of 5 CPE-like elements in a 3’UTR of 1269 nt in length.

Orb2 has essential functions during meiosis and subsequent differentiation of the spermatids. The 3’UTR deletion had no apparent effect on meiosis, and 64-cell spermatid cysts are formed without any visible defects. However, spermatid differentiation is disrupted. The earliest defect is observed in the polarization of the spermatid cyst. In WT cysts, the spermatid nuclei cluster towards the basal side of the cyst. This process is perturbed in *orb2*^*R*^, and the nuclei in a subset of mutant cysts remain scattered through the cyst. When *orb2*^*R*^ is combined with the null allele, *orb2*^*36*^, all cysts exhibit polarization defects. The 3’UTR deletion also disrupts the localization of *orb2* mRNAs and proteins near the tips of the elongating flagellar axonemes. Similar localization defects are observed for Boule and for *orb* mRNA and protein. While the progressive alterations in chromosome structure that accompany the maturation of the spermatids appear to take place, other steps in the maturation process are defective. These include the coalescence of the spermatid nuclei into a cap-like structure and the assembly and progression of the IC down the flagellar axoneme. Because of these defects, most *orb2*^*R*^ males are sterile, while the few that are fertile have a significantly reduced number of offspring.

Our results indicate that sequences in the *orb2* 3’UTR have important roles in spermatid differentiation, but they also raise several interesting questions. We have found that the levels of *orb2* transcript and protein in *orb2*^*R*^ are reduced about twofold. The simplest interpretation of this result is that the presence of intact 3’UTRs is important for *orb2* mRNA stability. However, in view of the sequence organization of the *orb2*^*R*^ deletion, it is also possible that the reduced amount mRNA and, consequently, protein levels is due to an inefficient use of the RH polyadenylation sequence. The approximately twofold reduction in the levels of *orb2*^*R*^ gene products is accompanied by variable and incompletely penetrant effects on spermatogenesis and male fertility. One explanation for these phenotypes is that *orb2* is haploinsufficient for several critical steps in spermatid differentiation and maturation. However, heterozygosity for *orb2*^*36*^ (an *orb2* deletion) results in a similar, if not greater, reduction in *orb2* transcripts and proteins without any concomitant effect on spermatogenesis or male fertility. Thus, a more likely explanation for the impairment of spermatogenesis and fertility in *orb2*^*R*^ is that the deleted 3’UTR sequences are required not only for normal mRNA accumulation but also for *orb2* function. Since *orb2* transcripts and proteins are not properly localized during flagellar axoneme elongation, a plausible conclusion is the deleted sequences are needed to facilitate the localization and/or translational regulation of *orb2* transcripts. In the deletion, insufficient amounts of Orb2 protein are produced in the cytoplasmic domains where it is required, and this, in turn, affects the localization of other transcripts and proteins (such as Boule and Orb) that have important roles in spermatogenesis.

Although our findings indicate that the deleted 3’UTR sequences are required for full *orb2* activity, the mutant phenotypes are variable and incompletely penetrant. This contrasts with the effects of deletions in the *orb* 3’UTR, which completely disrupts its key functions during oogenesis [42]. In the case of *orb2,* it is possible that there are other 3’UTR-independent mechanisms that can partially compensate for the defects in *orb2* function resulting from the 3’UTR deletion. It is also possible that this variability reflects the functional properties of the mutant *orb2* gene. In this respect, it is noteworthy that all transcripts produced by the *orb2*^*R*^ mutant, except RB are predicted to have a fairly long 3’UTR that retains five CPE-like elements. These 5 CPEs have sequences, UUUUUGU and UUUUUGUU, which were found to be enriched in Orb2-associated transcripts in tissue culture cells. Thus, the variable and incompletely penetrant phenotypes may arise because the 3’UTR still retains some functionality. Taken together, our data show that the 3’UTR of the *orb2* mRNA has a critical role in the regulation of its localization in spermatids. It implies the existence of *orb2* self-regulation feedback loop, which is important for male fertility.

## Materials and methods

### *Drosophila* stocks

The fly stock expressing Cas9 (#51324 from Bloomington Drosophila Stock Center) was used as WT control. The *orb2*^*36*^ stock (#58479 in the Bloomington Drosophila Stock Center) was described previously [18].

### Generation of *orb2*^*R*^ allele

Two guide RNAs were used to delete the portion of *orb2* 3’UTR using the CRISPR/Cas9 system (see Supplement). gRNA sequences were cloned into pU6 vector (pU6-gRNA, Addgene plasmid # 5306; a gift from Caixia Gao), and corresponding 1-kbp homology arms were cloned into pHD vector (pHD-DsRed, Addgene plasmid #51434; a gift from Kate O’Connor-Giles). These constructs were injected into #51324 fly stock, and dsRed-positive flies were selected. The marker was removed using loxP sites; and the 3’UTR of *orb2*^*R*^ was sequenced.

### Fertility assay

Groups of 20–28 male flies of the WT, *orb2*^*R*^ and *orb2*^*R*^/*orb2*^*36*^ genotypes were individually crossed with two WT virgin females for 7 days, and then adult flies were removed from the vials. The presence of larvae, pupae, and adults in the vials was examined after another 2 weeks. The males that were able to mate and produce larvae were regarded as fertile.

### Breeding efficiency analysis

A total of 175 males of the WT, *orb2*^*R*^ and *orb2*^*R*^/*orb2*^*36*^ genotypes were individually crossed to WT virgin female flies for 7 days. The adult flies were then removed, and the numbers of offspring from each individual male were estimated and ranged into groups.

### Viability assay

Viability test was based on Mendelian inheritance in the offspring of *orb2*^*R*^ and *orb2*^*R*^/*orb2*^*36*^ alleles. For *orb2*^*R*^, individual *orb2*^*R*^/*TM3 Ser* male and female flies were crossed with each other for 7 days; for *orb2*^*R*^/*orb2*^*36*^, crosses were made between individual *orb2*^*36*^/*TM3 Ser, Sb* males and *orb2*^*R*^/*TM3 Ser* females. After the next 2 weeks, the phenotypic ratio in the offspring was evaluated by chi-square analysis.

### Antibodies

The antibodies used were as follows: mouse anti-Orb2 (4G8) at 1:100 for Western blotting, mouse anti-Orb2 (4G8 & 2D11) at 1:25 and mouse anti-Orb (6H4) at 1:30 for whole testis staining. These antibodies were produced and deposited to the DSHB by P. Schedl. Rabbit anti-bNactes (used at 1:300) was a gift of Dr. G.L. Kogan (Institute of Molecular Genetics). Rabbit anti-Bol (used at 1:1500) was a gift from Steven Wasserman. Secondary antibodies were goat anti-mouse IgG conjugated with Alexa 488 or 546 and goat anti-rabbit IgG conjugated with Alexa 546 (Invitrogen). Alexa 633–phalloidin at 1:300 (Thermo Fisher Scientific) was used for actin staining in whole mounts of testes.

### Whole mount immunostaining

Testes from 1-to 3-day males were dissected in PBST (0.1% Tween-20 in 1× PBS), fixed in 4% paraformaldehyde for 20 min, washed with three portions of PBST (here and below, each wash for 5 min), and then passed through an ascending-descending methanol wash series (30%, 50%, 70%, 100%, 70%, 50%, 30% in 1×PBS). The testes were then washed with two portions of PBST and incubated in PBSTX (0.1% Tween-20 and 0.3% Triton X-100 in 1×PBS) with 5% normal goat serum (Life Technologies) at room temperature for at least 1 h. This was followed by overnight incubation with primary antibody and, after washing with three portions of PBSTX, with secondary antibody at room temperature for at least 2 h. After final washing with three portions of PBSTX, the preparations were mounted on slides in VECTASHIELD mounting medium with DAPI (Vector Laboratories).

### Fluorescence in situ hybridization

Quasar 670-conjugated *orb2* and *orb* FISH probes were from LGC Biosearch Technologies [27]. Testes were taken from young male flies that were fed yeast paste for 2–3 days. They were dissected in 1×PBS, fixed in 4% paraformaldehyde for 30 min, rinsed in four portions of PBST, dehydrated through an ascending methanol series, and stored in 100% methanol at −20⁰C for 10 min. After rehydration in PBST, the testes were additionally rinsed in four portions of PBST, transferred to wash buffer (4×SSC, 35% formamide, 0.1% Tween-20) for 15 min at 37⁰C. This was followed by incubation with the FISH probes overnight at 37⁰C in hybridization buffer (10% dextran sulfate, 0.01% salmon sperm single-strand DNA, 1% vanadyl ribonucleoside, 0.2% BSA, 4×SSC, 0.1% Tween-20, and 35% formamide). The resulting preparations were washed in two portions of wash buffer, 1 h each, at 37⁰C and mounted in Aqua-Poly/Mount (Polysciences, Inc.).

### Microscopy

Stained preparations were scanned and imaged under an LSM 510 META confocal laser scanning microscope (Carl Zeiss Jena, Germany) in multichannel mode using 63× or 40× oil objective lenses and 10 × air objective lens (numerical aperture 1.4). Images with a frame size of 1024 × 1024 pixels and a z resolution of 1 μm were taken at a scan speed of 7, in four replicates, and imported into Imaris 5.0.1 (Bitplane) and Adobe Photoshop for subsequent processing.

### RNA isolation, reverse transcription, and qPCR

Testes of 1-to 3-day male flies were dissected in cold 1× PBS. Total RNA was isolated from 25 pair of testes using TRIzol (Life Technologies) according to the manufacturer’s protocol, treated with DNase (TURBO DNA-free kit, Thermo Fisher Scientific), and reverse transcribed into cDNA. The level of the transcripts was estimated using gene-specific primers (see Supplement). RT–qPCR for each sample was performed in technical triplicate. The data presented correspond to the mean of 2^−ΔΔCt^ from at least ten independent experiments.

### Semi-quantitative western blot

Testes of 1-3 day male flies were dissected in cold 1× PBS and immediately transferred to lysis buffer (100 mM KCl, 5 mM MgCl_2_, 10 mM HEPES, 0.5% NP-40, 1 mM DDT, PIC, PMSF). Total protein lysates were prepared from 25 pair of testes and loaded in equal dilution series (5, 10, and 20 μg) onto precast stain-free PAAG gel (Bio-Rad). After electrophoresis, the gel was visualized in a ChemiDoc system (Bio-Rad) to evaluate protein concentrations and perform normalization against the total protein level. The proteins from the gel were blotted onto a PVDF membrane, which was incubated with primary antibodies, secondary HRP-conjugated antibodies, and the Super Signal Western Femto substrate (Thermo Fisher Scientific). The induced chemiluminescence was measured with a ChemiDoc visualization system.

### Quantification and statistical analysis

The *orb2* mRNA and protein enrichment was calculated using average intensity projections of the growing end of spermatid cysts (Fig. 4C). First, the mean fluorescence intensity of the growing tip of the flagellar axoneme was determined by averaging several z-stacks in the three areas of interest. The mean fluorescence intensity in spermatocytes was determined in the same way. Then the mean fluorescence intensity of each area of interest within a spermatid was divided by the mean fluorescence intensity of spermatocytes for each testis. These ratios are shown as box plots. The Imaris software was used to quantify the fluorescence signal of *orb2* mRNA and Orb2 protein.

Experimental data were processed statistically with the GraphPad Prism software. The statistical significance of the observed differences was estimated by unpaired two-tailed *t*-test (Figs. 3A, 4C). Mendelian inheritance in the offspring was analyzed using the nonparametric chi-square method. Variable values for each group are presented as the mean ± standard deviation (SD) or ± standard error of mean (SEM). For all panels, \**P* < 0.05, \*\**P* < 0.005, \*\*\**P* < 0.0005, \*\*\*\**P* < 0.0001; ns, not significant.

## Acknowledgments

This study was supported by the Russian Science Foundation (grant 18-74-10051 to MZ) and a NIH (R35GM126975 to PS). The authors are grateful to the Center for Precision Genome Editing and Genetic Technologies for Biomedicine of IGB RAS for mRNA and protein quantification, the Core Facilities Center of IGB RAS, and the IMG RAS Core Facility “Center of Cell and Gene Technologies” for providing the equipment for microscopy.

## Supplementary Information

**Suppl. Fig 1.**
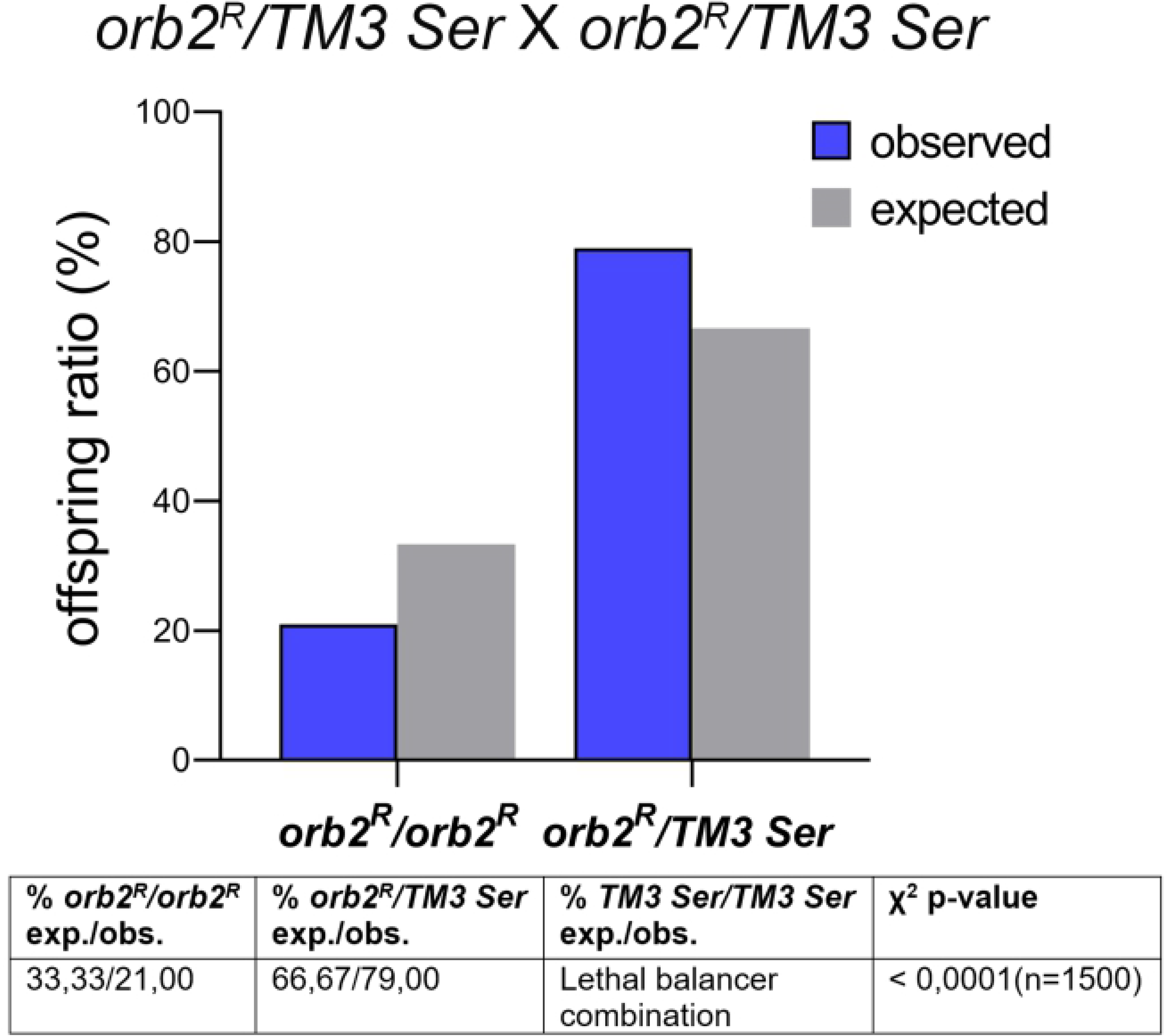
Mendelian inheritance of *orb2*^*R*^ allele. Observed and expected offspring ratios and chi-square analysis of offspring with different genotypes from *orb2*^*R*^/*TM3 Ser* intercross.

**Suppl. Fig 2.**
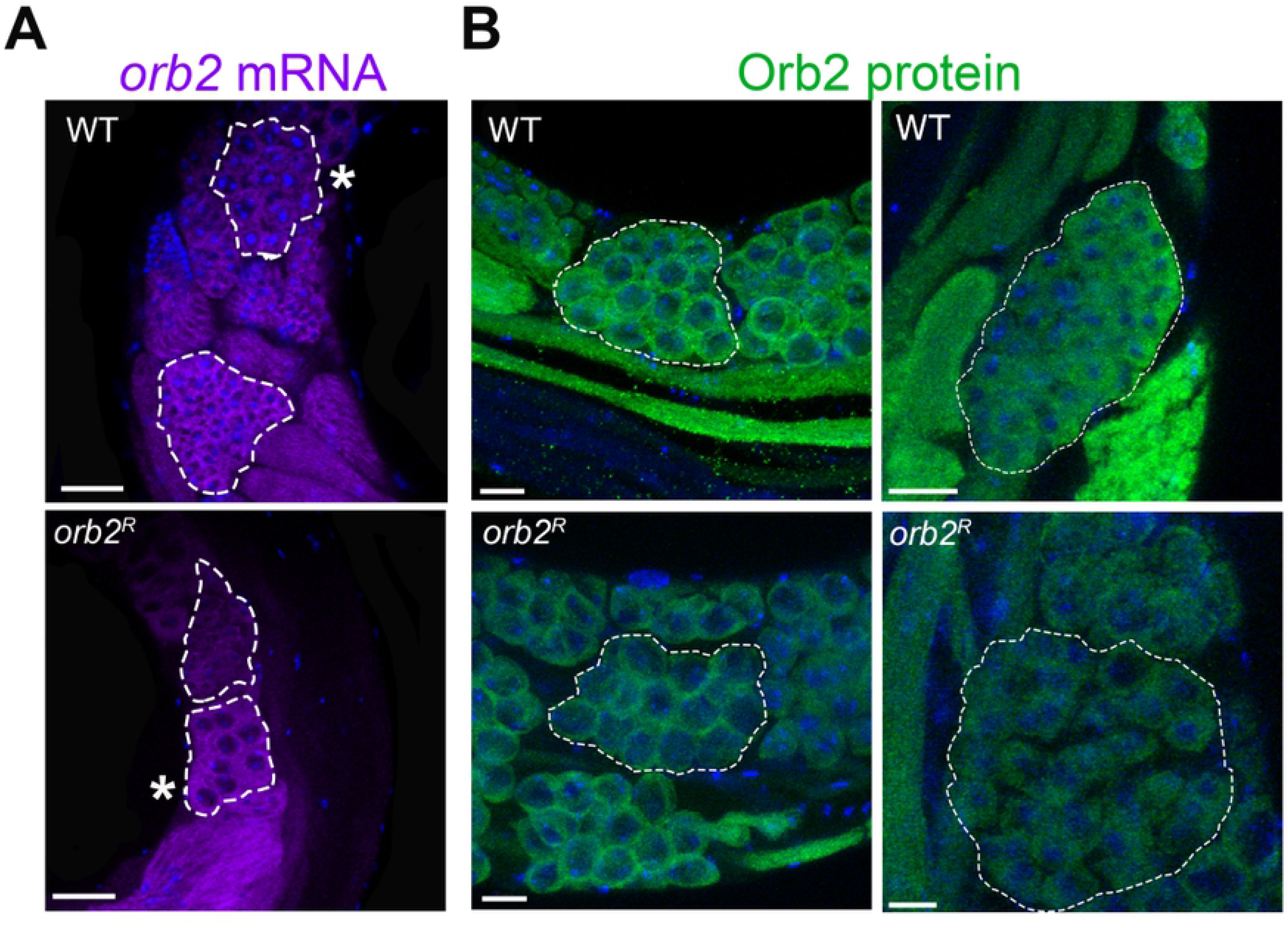
Orb2 protein and *orb2* mRNA in premeiotic and meiotic cysts. **(A)** Fluorescence in situ hybridization for *orb2* mRNA localization in primary spermatocytes (16-cell premeiotic cysts) within the area indicated by an asterisk and in secondary spermatocytes (32-cell meiotic cyst) within the highlighted area without an asterisk in WT and *orb2*^*R*^ testes. Except for a slight decrease in the signal, the mutants show no obvious changes in the localization of *orb2* mRNA at these stages (WT, *n* = 25; *orb2*^*R*^, *n* = 18). **(B)** Whole mount staining of testes with Orb2 antibodies shows that the level of Orb2 protein in primary spermatocytes (left column) in *orb2*^*R*^ is reduced, compared to WT, with the pattern of its localization remaining unchanged; the same is also true of secondary spermatocytes (right column) (WT, *n* = 20; *orb2*^*R*^, *n* = 21). Scale bar, 30 μm.

**Sequences for Cas9 cut**

upstream cut, 5’- CTTCGTAATAGACCGTATTAT↓GTAAGG;

downstream cut, 5’- CTTCGTGGAGTACTGCTGATA↓TCTTGG.

**Primers used in qPCR**

*orb2* total mRNA, 5’-TAACACCAGCGAAAGGGGAC and 5’- CAGATGTGCGACGAGTGC;

*orb2* primary transcript, 5’-GCTGTTGGTGCTGATGGA and 5’-AGCCTCTTCATCTTGTTGTC;

*GAPDH* mRNA, 5’-CTACCTGTTCAAGTTCGATTCGAC and 5’- AGTGGACTCCACGATGTATTCG.

